# Seed mass and plant origin interact to determine species germination patterns

**DOI:** 10.1101/841114

**Authors:** Andrea Veselá, Tomáš Dostálek, Maan Rokaya, Zuzana Münzbergová

## Abstract

Ongoing changes in temperature and precipitation regime may have strong impact on vulnerable life-history stages such as germination. Differences in germination patterns among species and populations may reflect their adaptation to conditions of their origin or may be determined by the phylogenetic constrains. These two effects are, however, rarely separated. All the germination patterns may also be modified by seed mass.

We studied 40 populations of 14 species of *Impatiens* coming from Himalayas. Germination of seeds of different origin was tested in four target temperatures, three simulating original conditions plus a warmer climate change scenario. We also studied effect of shorter stratification and warmer temperature in combination as another possible effect of climate change.

Original and target climate interacted and had strong impact on total germination, but not on germination speed and seed dormancy. Interaction between seed mass and original climate indicated different germination strategies in light and heavy seeds. Only seed mass was affected by phylogenetic relationships among the species, while germination response (with exception of T50) was driven primarily by climate of origin.

This study is the first to show that the effect of seed mass interacts with original climate in determining species germination patterns under changing climate. The differences in seed mass are thus likely crucial for species ability to adapt to novel conditions as seed mass, unlike seed germination patterns, is strongly phylogenetically constrained. Further studies exploring how seed mass modifies species germination under changing climate are needed to confirm generality of these findings.

## Introduction

Ongoing as well as forecasted increase in temperature and changes in precipitation regime (IPCC 2014) may have strong impact on vulnerable plant life-history stages such as germination, seedling emergence and seed production (Knapp et al. 2008) (Walck et al. 2011) (Dreesen et al. 2014) due to their strong dependency on temperature and moisture (Baskin and Baskin 2001, Fenner and Thompson 2005). Generally, it is known that original (Meyer, Allen and Beckstead 1997) (Santo, Mattana and Baechetta 2015) (Seglias et al. 2018) as well as target (e.g. (Grime et al. 1981) (Schütz and Rave 1999) (Gardarin, Daurr and Colbach 2011) environment affects germination at both interspecific (e.g. (Gimenez-Benavides, Escudero and Perez-Garcia 2005) and intraspecific level (e.g. (Qaderi and Cavers 2002) (Tingstad et al. 2016). However, only a few studies focused on the interactions of original and target climate (reviewed in (Walck et al. 2011)) on species germination though these interactions are known to strongly affect plant performance (Münzbergová et al. 2017) as well as species germination (Veselá et al. submitted).

In nature, species are exposed to cold stratification, which is often necessary for dormancy breaking especially in mountain (Meyer 1992) and temperate zone species (Baskin and Baskin 1988). Even though some species are able to germinate without stratification, species with stratification germinate more and faster e.g. (Martin 1965) (Cavieres and Arroyo 2000) (Garcia-Fernandez et al. 2015). Also, the duration of stratification plays an important role in species germination. Some species germinate well after one or two months of stratification (Carta et al. 2014) (Perglová et al. 2009), while other require stratification of six and more months (Schütz and Milberg 1997) (Esmaeili et al. 2009) (Perglová et al. 2009) (Garcia-Fernandez et al. 2015). Necessary stratification duration can vary among species but also among populations of one species, as demonstrated by Cavieres and Arroyo (2000). Seeds from higher altitudes require longer stratification than seeds from lower altitudes. The ongoing climate change may shorten the stratification period (IPCC 2014) possibly leading to decreasing germination in some species (Garcia-Fernandez et al. 2015) (Carta et al. 2016b). However, the effects of reduced stratification on species germination in warmer climates have not yet been studied.

Germination characteristics do not depend only on environmental conditions but also on seed traits such as seed mass (Wang et al. 2009) (Wang et al. 2012) (Liu et al. 2013) (Hradilová et al. 2019). (Rees et al. 2001) demonstrated that heavier seeds have higher content of nutrients and higher germination percentage. Positive effect of seed mass on germination was also confirmed in a study of (Navarro and Guitian 2003) and (Münzbergová and Plačková 2010) at intraspecific level and (Wu and Du 2007), and (Paulů, Harčariková and Münzbergová 2017) at interspecific level. However, some studies showed an opposite trend (e.g. (Wang et al. 2009) (Wu, Li and Du 2011)). The effects of seed mass on species response to changing climatic conditions, however, remain to be explored.

Seed traits including seed mass are strongly associated with phylogeny (Zhang, Du and Chen 2004) (Moles et al. 2005) (Norden et al. 2009) (Barak et al. 2018). Phylogeny explains considerable part of variance in germination among species (Bu et al. 2008) (Wang et al. 2009) (Xu et al. 2014) (Seglias et al. 2018) and strong phylogenetic signal was found also in seed dormancy traits (Dayrell et al. 2017). The likely cause is that phylogeny imposes limits to variability in reproductive attributes within a clade because of similar developmental and design constraints in related species (Ackerly and Donoghue 1995) (Figueroa and Armesto 2001). However, to our knowledge, there is only one study focusing on the effects of within genus phylogeny on species germination. It is based on seven species and showed that more closely related species had more similar germination behavior and the authors point out that this subject requires further attention (Carta, Hanson and Muller 2016a). Such knowledge is likely to increase our understanding of evolution of species germination requirements.

In this study, we explored germination response and necessary stratification duration of 40 populations belonging to 14 species of genus *Impatiens* coming from the Himalaya mountains in Nepal. The aim of our study was to answer the following questions: i) What is the effect of seed mass and original and target conditions on species germination and the necessary stratification duration?, ii) Does shorter stratification influence germination response to warmer conditions?, iii) Are the patterns affected by phylogenetic relationships among the species?

We hypothesized that germination will increase with increasing seed mass. It will also increase with increasing original as well as target temperature and the original and target effects will interact with each other. Populations from warmer conditions will require shorter stratification than populations from colder conditions and reduced stratification will reduce germination in warmer temperature. Simultaneously, we predict that more closely related species will have more similar germination response and the effects of seed mass and target and original conditions on seed germination will be thus modified when accounting for species phylogeny.

## Methods

### Study species and seed collection

We used species of the genus *Impatiens*, Balsaminaceae, for the study. Genus *Impatiens* is highly diversified genus of annual or perennial herbs comprising over 1 000 species, generally occurring at high altitudes, i.e., more than 1 500 m above sea level, distributed in the mountains of the Old World tropics and subtropics, with only few species occurring in northern hemisphere temperate regions (Grey-Wilson 1980) (Yuan et al. 2004) (Janssens et al. 2009). One of the biodiversity hotspots of the genus is found in eastern Himalayas and south-east Asia (Song, Yuan and Kupfer 2003) (Yuan et al. 2004) (Yu et al. 2016), i.e. the region of our study. The species occupy diverse habitats such as forest understory, roadside ditches, valleys, abandoned fields, stream banks and seepages (Yu et al. 2016).

In total, we used seeds collected from 40 populations, with seeds from 24 populations collected in 2016 and 16 populations in 2017 belonging to 14 species (list of species and locations of their collection are provided in Supporting information 1). Seeds in each population were collected from at least 5 maternal plants. Seeds from each maternal plant were kept separately during storage and germination tests. After collection, the seeds were stored at ambient moisture (approximately 55%) at room temperature (approximately 18 °C) for 5 months before the beginning of the experiments. For each population, we recorded year of collection and origin of the population (altitude, longitude, latitude) and determined weight of one thousand seeds. Further we determined original temperature and original precipitation, which were derived from WorldClim database (Hijmans et al. 2005) according to the coordinates of the populations. We used mean temperatures from March to June since it represents premonsoon period when most *Impatiens* species germinate and start to grow.

### Germination tests

Undamaged, fully developed, seeds were used for the germination tests. For each population and growth chamber (see below), we established 5 replicates (each replicate corresponding to one mother plant) with 12 seeds each. The seeds were incubated on moist filter paper in 5 cm diameter Petri dishes. In cases, when we had deficiency of seeds, we used less than 12 seeds per Petri dish but never less than 6. First, the seeds on Petri dishes were exposed to cold stratification (5°C). When at least one seed on 30% of Petri dishes germinated, all Petri dishes of that population were transferred to growth chambers (describes below) and the time was recorded as the necessary stratification duration. The germinated seeds were recorded and removed every week. The seed was considered germinated if the radicle was visible to the naked eye. Weekly, we also removed all the rotten seeds from the dishes. When no seed of any population germinated for three subsequent weeks, the experiment was terminated. Healthy-looking ungerminated seeds were tested for viability by tetrazolium chloride method according to (Cottrell 1947).

We used four combinations of germination conditions in the growth chambers, further referred to as target conditions. The target conditions were set up with respect to the course of temperatures during the day on the original localities: 1) the coldest regime (further referred to as 6/17.5°C on the basis of minimum and maximum temperatures) - mean temperature from March to June in 2 700 m a.s.l., i.e. in the altitude representing median of upper altitudinal limits of *Impatiens* species in Nepal, 2) cold regime (further referred to as 9/20°C) - mean temperature from March to June in 2 250 m a.s.l., i.e. in the altitude representing median of centres of altitudinal ranges of *Impatiens* species in Nepal, 3) warm regime (further referred to as 12/22.5°C) - mean temperature from March to June in 1800 m a.s.l., i.e. in the altitude representing median of the lowest altitudinal limits of *Impatiens* species in Nepal, and 4) the warmest regime (further referred to as 15/25°C) - mean temperature from March to June in 1 800 m a.s.l., i.e. in the altitude representing median of the lowest altitudinal range of *Impatiens* species in Nepal, in year 2050 as predicted by global climate model MIRO5C at RCP8.5 (Tatebe et al. 2012).

Information on altitudinal ranges of *Impatiens* species in Nepal was obtained from Annotated Checklist of the Flowering Plants of Nepal (http://www.efloras.org/flora_page.aspx?flora_id=110), which is an updated online version of (Press et al. 2000). Temperature data were obtained from WorldClim database (Hijmans et al. 2005). Data on mean temperatures in particular altitudes were obtained from slopes of correlations between altitudes and mean temperatures for particular data points along four valleys in Central and East Nepal where our seed collections took place. We used mean temperatures from March to June since it represents premonsoon period (see above). The course of the temperatures during the day was modelled based on mean, minimum and maximum temperatures. The lowest day temperatures were from 3 to 5 am, the highest day temperatures were from 12 to 14 pm, for all details see Supporting information 2. For all the regimes, the same day length and radiation were used, i.e. 12 h of 60% light (06.00–18.00 h; 250 μmol m^−2^ s^−1^) and 10 h of full dark with a gradual change in light availability in the transition between the light and dark period over 1 h. Petri dishes were regularly watered with demineralized water.

For seeds collected in 2017, only three growth chambers were available for our experiments due to technical constrains. For these seeds we thus did not use the 9/20°C regime. Instead, we added one additional germination regime with all seeds only stratified for one month and then transferred to the growth chamber with the highest temperature (15/25°C). This temperature sequence simulated possible impact of climate change, i.e. shortening of stratification and warmer climate. This regime was not used in the main statistical analyses presented in Table 1 and was only compared with the corresponding 15/25°C regime with longer stratification period (see above).

**Table 1.** Effect of original temperature, original precipitation, seed mass and target temperature on total germination, seed dormancy, germination speed (T50) and necessary stratification duration assessed using generalized linear mixed effects models with population used as a random factor. Year, longitude, latitude and their interaction were used as covariates in the tests. N.t. indicates not tested. Significant values (≤ 0.05) are in bold. * indicates results becoming non-significant after including phylogeny.

|  | Total germination |  | T50 |  | Seed dormancy |  | Necessary stratification duration |  |
| --- | --- | --- | --- | --- | --- | --- | --- | --- |
|  | F-value | p-value | F-value | p-value | F-value | p-value | F-value | p-value |
| Orig.temp | 0.63 | 0.673 | 2.47 | 0.083 | 0.01 | 0.899 | **4.94** | **0.033** |
| Orig. precip | 0.58 | 0.698 | 1.73 | 0.182 | 0.60 | 0.701 | **13.85** | **0.001** |
| Target temp | **5.98** | **0.015** | **17.14** | **<0.001\*** | **3.76** | **0.049** | N.t. |  |
| Seed mass | **11.55** | **<0.001** | **16.56** | **<0.001\*** | 0.27 | 0.860 | 0.75 | 0.387 |
| Orig.temp: Orig.precip | 0.11 | 0.729 | 1.47 | 0.179 | 1.41 | 0.171 | **12.28** | **0.001** |
| Orig.temp: Target temp | 0.39 | 0.239 | 2.10 | 0.209 | 0.24 | 0.682 | N.t. |  |
| Orig.precip: Target temp | **4.04** | **0.035** | **9.53** | **0.014\*** | 1.26 | 0.235 | N.t. |  |
| Orig.temp: Seed mass | **13.43** | **<0.001** | 0.73 | 0.559 | **3.74** | **0.042** | 0.30 | 0.585 |
| Orig.precip: Seed mass | **17.50** | **<0.001** | 2.20 | 0.153 | **6.44** | **0.018** | **5.94** | **0.015** |
| Target temp: Seed mass | 0.22 | 0.632 | 2.36 | 0.119 | 1.52 | 0.274 | N.t. |  |
| Orig.temp: Orig.precip: Target temp | **13.72** | **<0.001** | 2.98 | 0.070 | 2.26 | 0.121 | N.t. |  |
| Orig.temp: Orig.precip: Seed mass | **4.64** | **0.024** | **29.61** | **<0.001\*** | 0.04 | 0.896 | **4.35** | **0.038** |
| Orig.temp: Target temp: Seed mass | 2.08 | 0.106 | 0.09 | 0.902 | 1.62 | 0.159 | N.t. |  |
| Orig.precip: Target temp: Seed mass | 0.95 | 0.297 | 0.19 | 0.635 | 2.97 | 0.070 | N.t. |  |
| Orig.temp: Orig.precip: Target temp: Seed mass | 0.01 | 0.935 | 3.07 | 0.069 | 3.06 | 0.051 | N.t. |  |

### Data analysis

The dependent variables in our analyses were total germination, time to 50% germination (T50), germination index (GI – describes ratio of the germination percentage and speed), seed dormancy, seed viability and necessary stratification duration. Total germination was expressed as the proportion of germinated seeds from all seeds in one Petri dish over the whole period. T50 was calculated according to (Coolbear, Francis and Grierson 1984) modified by (Farooq et al. 2005). Germination index was calculated as described in the Association of Official Seed Analysts (AOSA 1983). GI resp. T50 cannot be calculated when 0 resp. 0 or 100% seeds of all seeds in Petri dish germinate and the corresponding cases were thus excluded from this calculation. Seed dormancy was defined as the proportion of seeds, which were found to be viable after tetrazolium chloride application. Seed viability was sum of proportion of germinated and dormant seeds. Necessary stratification duration is described above (time when at least one seed on 30% of Petri dishes germinated). Pair-wise correlation matrix of the variables, based on Pearson’s correlation coefficient, is presented in Supporting information 3. Because of strong correlation of GI and seed viability with total germination and dormant seeds (r ≥ 0.577), we did not use the seed GI and seed viability in further analyses.

We tested the effect of target temperature (temperature in the growth chambers), original temperature, original precipitation, seed mass and all their interactions on all the dependent variables. To assess whether seed mass is affected by original climate, we tested the effect of original temperature, original precipitation and their interaction on seed mass. All the tests were done using generalized linear mixed effect models as implemented in the lme4 package in R (Bates et al. 2015) with population as a random factor. We assumed binomial distribution of total germination and seed dormancy (information on number of germinating/dormant and non-germinating/non-dormant seeds linked using c-bind function in R). T50 and necessary stratification duration followed Gaussian distribution after log-transformation. No transformation was necessary for seed mass. In all analyses, year of seed collection, longitude, latitude and interaction of longitude and latitude were used as covariates.

Because for seeds collected in 2017, we did not study the germination in one of the target temperatures (9/20°C) due to technical constrains, we repeated all the tests after excluding this regime from the total dataset. The results of these analyses are presented only in Supporting information 4, as they were in most cases in line with analyses including data from all the target temperatures.

To assess whether the combination of shorter stratification and warmer temperature has any effect on seed germination response, we conducted a separate comparison of seeds exposed to shorter stratification and warmer temperature and seeds kept under stratification until 30% germination and warmer temperature. We used generalized linear mixed effect models with population as a random factor, as described above, for these analyses. Effect on number of dormant seeds was not tested, as no dormant seeds were found in the warmer regime (i.e. in 15/25°C).

To assess the effect of phylogenetic relationships among the species on the patterns observed, we used ITS based phylogeny of the plant group developed for the purpose of another study (Líblová et al, in prep.). Phylogenetic distance matrix describing the relationships among species was decomposed into its eigenvectors using PCoA as suggested by Diniz et al. (1998) and Desdevises et al. (2003) using the R-package ‘ape’ (Paradis et al. 2004). The first two eigenvectors explained 76 % of the variability in the data. Their effect was tested on all dependent variables and seed mass. In cases, where eigenvectors had significant effect, they were included as co-variables in the above described models in order to correct for phylogenetic autocorrelation and to compare the effects of phylogeny to the effects of original and target environment.

## Results

### Effect of original and target environment and seed mass

Seed mass was not significantly affected by original temperature, original precipitation or their interaction (all p-values ≥ 0.707). All the germination characteristics were independent of original temperature, original precipitation and their interaction with exception of necessary stratification duration (Table 1). Necessary stratification duration increased with increasing original temperature. The effect of original temperature on the necessary stratification duration also interacted with original precipitation, with the necessary stratification duration being the highest in seeds coming from warm and dry localities and the lowest in seeds coming from cold and wet localities (Fig 2).

All the germination characteristics were affected by target temperature (Table 1), but with low explanatory power (Fig 1). T50 increased and total germination and seed dormancy decreased with increasing target temperature. Total germination and T50 were also significantly affected by seed mass (Table 1) with heavier seeds germinating more but slower than lighter seeds. The explanatory power of seed mass was similar to that of target temperature (Fig 1).

**Fig 1.**
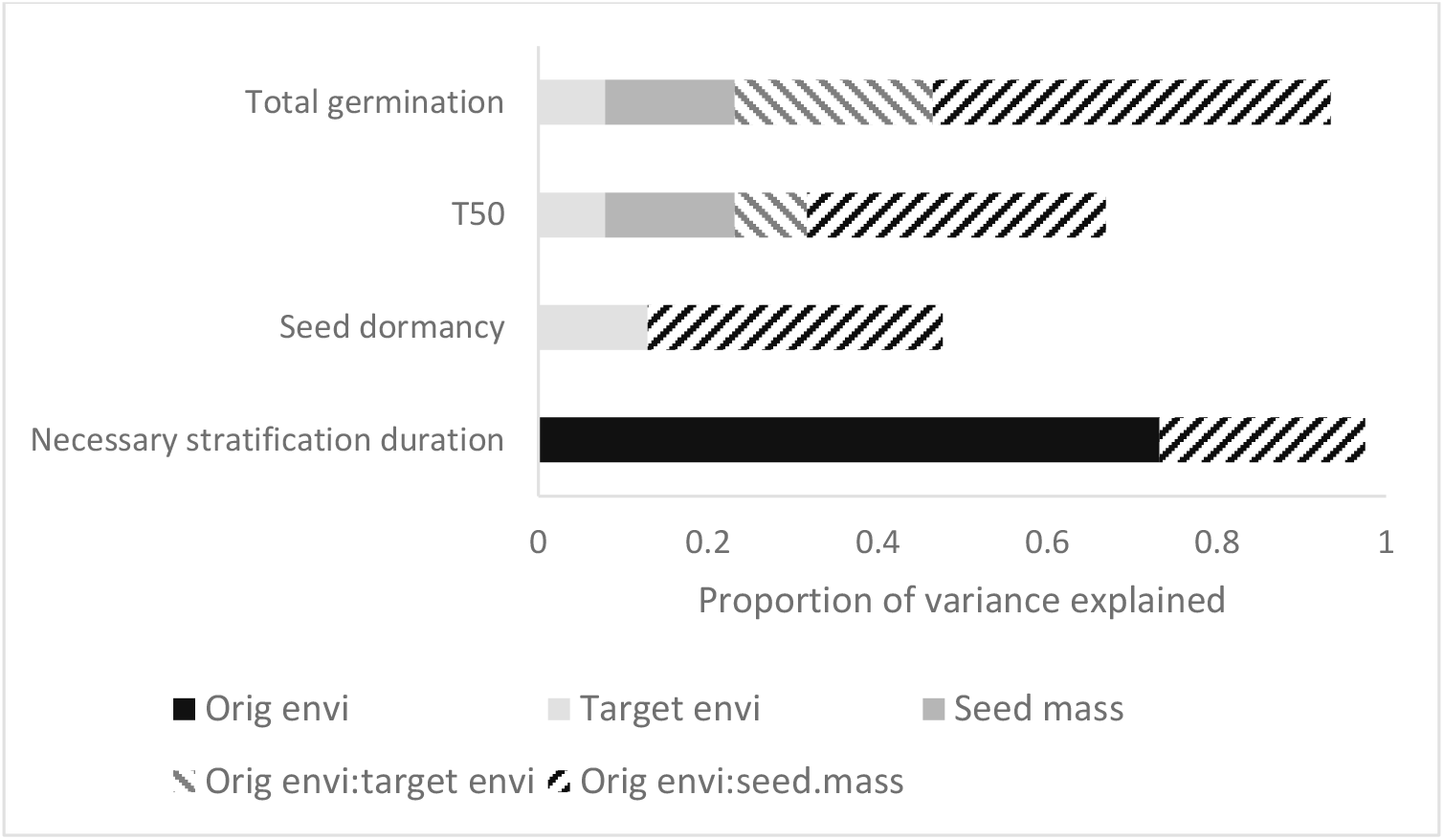
Proportion of variance explained by original environment (including both temperature and precipitation), target temperature, seed mass and their interaction on total germination, germination speed (T50), seed dormancy and necessary stratification duration. Interaction of target environment and seeds mass and triple interaction of original environment, target environment and seed mass are not figured in the graph as proportion of variance explained was very low.

**Fig 2.**
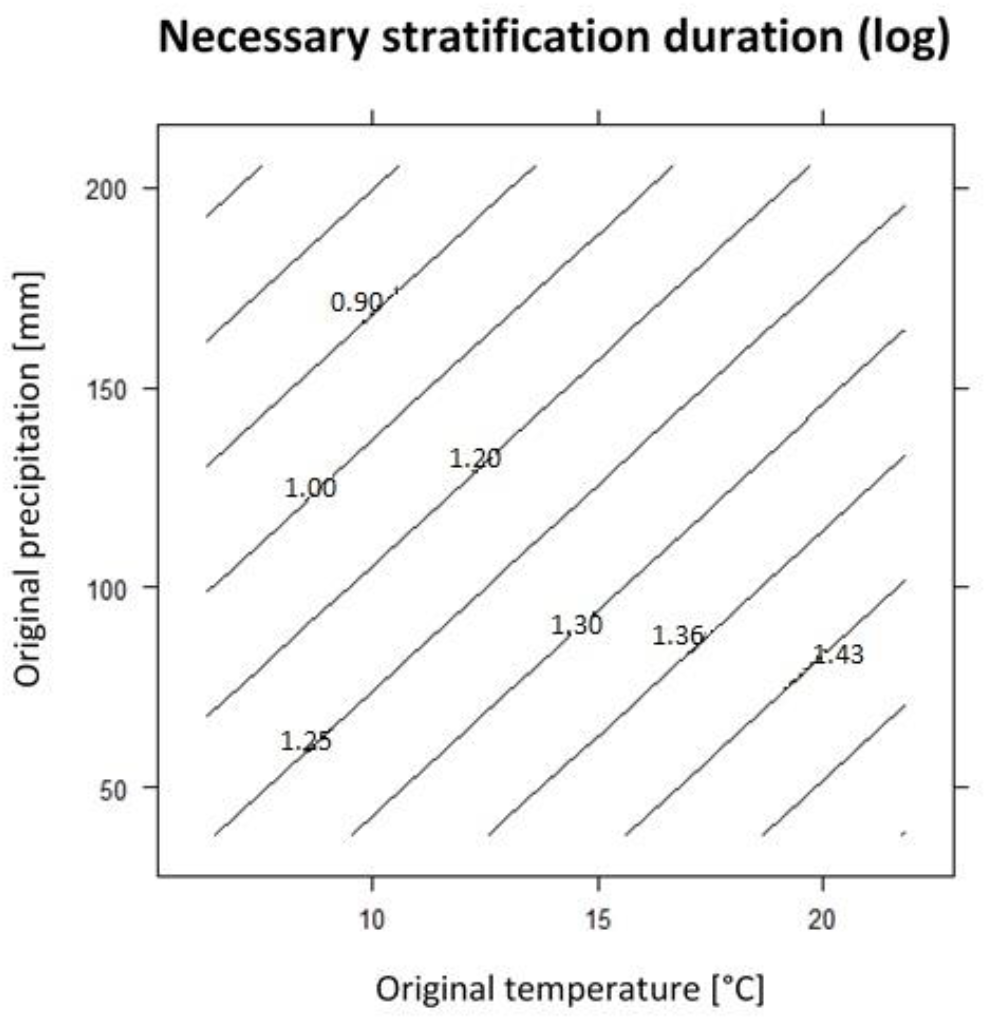
Effect of original temperature and original precipitation on necessary stratification duration in weeks (log).

Total germination and T50 were also affected by the interaction between original precipitation and target temperature (Table 1). Total germination was the highest in seeds coming from the wettest localities exposed to 9/20 °C and the lowest in seed coming from the driest localities exposed to 12/22.5 °C. In contrast, T50 was the highest in seeds coming from the wettest localities exposed to 6/17.5 °C and the lowest in seed coming from the wettest localities exposed to 15/25 °C.

Seed mass significantly interacted with original temperature and also with original precipitation in its effects on total germination, seed dormancy and necessary stratification duration (Table 1) with high proportion of variance explained in total germination and seed dormancy (Fig 1). Total germination and seed dormancy were the highest in heavy seeds coming from the warmest localities and the lowest in light seeds coming from the coldest localities. Total germination was the highest in heavy seeds coming from the driest localities and the lowest in light seeds coming from the wettest localities and the germination was more dependent on seed mass than on original precipitation. Seed dormancy was the highest in heavy seeds coming from the warmest localities and the lowest in light seed coming from the coldest localities. Necessary stratification duration increased with increasing seed mass and original precipitation.

Total germination was also affected by triple interaction of original temperature, original precipitation and target temperature with quite high proportion of variance explained (Fig 1). In all target temperatures with exception of 12/22.5 °C, total germination increased with increasing original temperature and decreasing original precipitation. In 12/22.5 °C, germination increased with decreasing original precipitation and original temperature. Total germination in these target conditions was more dependent on original precipitation than on original temperature, while in the three other target conditions germination was affected equally by both variables. The highest germination was observed in the coldest target conditions, i.e. 6/17.5 °C (Fig 3).

**Fig 3.**
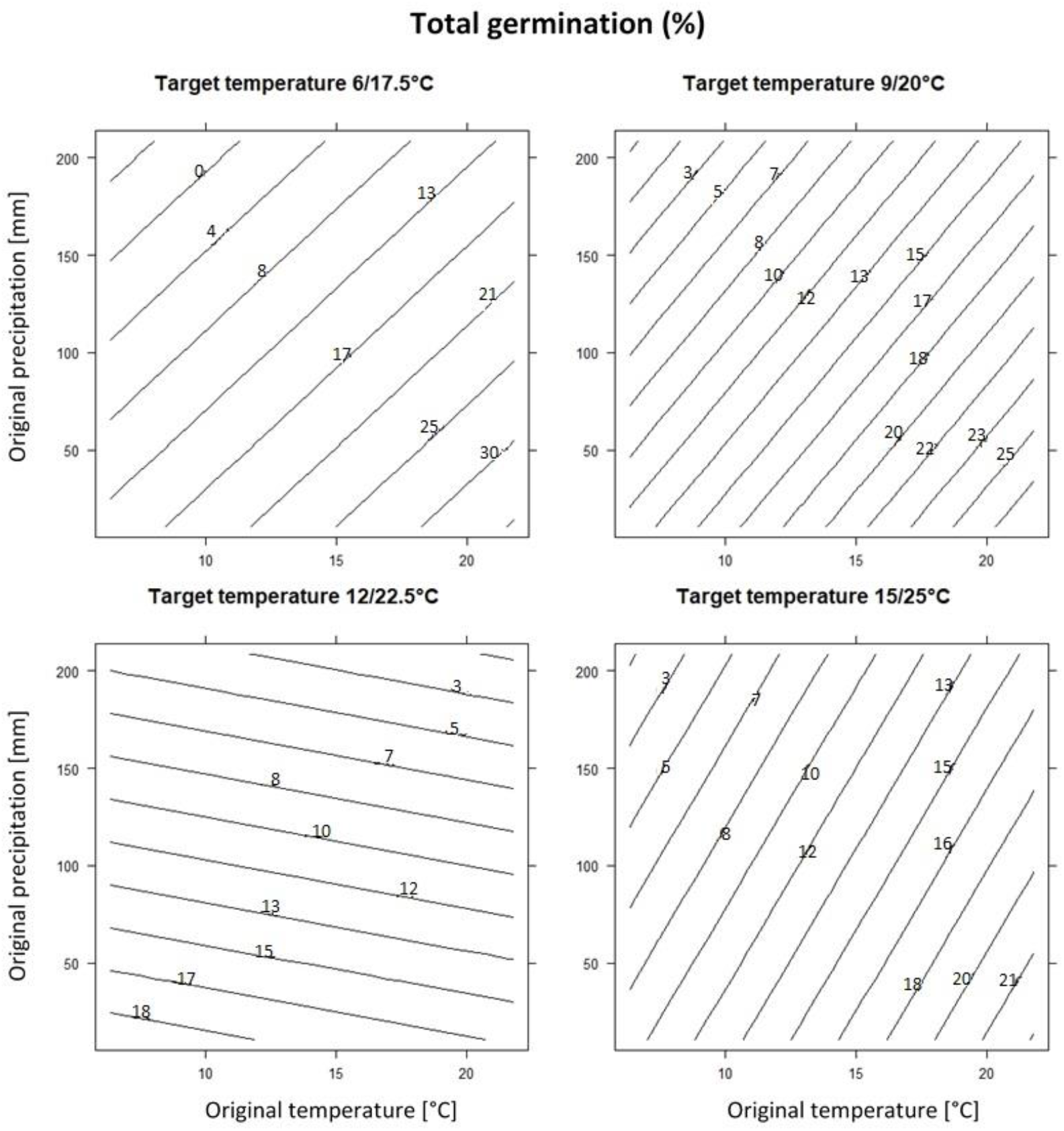
Effect of target temperature, original temperate and original precipitation on total germination (%).

Total germination and T50 were significantly affected by the interaction of original temperature, original precipitation and seed mass (Table 1) with high explanatory power both in total germination and in T50 (Fig 1). Lighter seeds coming from colder and wetter localities germinated in high number and quite fast. In contrast, heavier seeds germinated in high numbers, when they came from warmer and drier localities. Fast germination was in seeds coming from warmer and wetter localities and was more dependent on original temperature than on original precipitation (Fig 4). Significant triple interaction between seed mass, original temperature and original precipitation (Table 1) also indicated that the stratification duration increased in lighter seeds with decreasing original temperature and original precipitation and the stratification duration was more dependent on original precipitation than on original temperature. The necessary stratification duration in heavy seeds increased with increasing original temperature and decreasing original precipitation and the stratification duration was more dependent on original temperature than on original precipitation (Fig 5).

**Fig 4.**
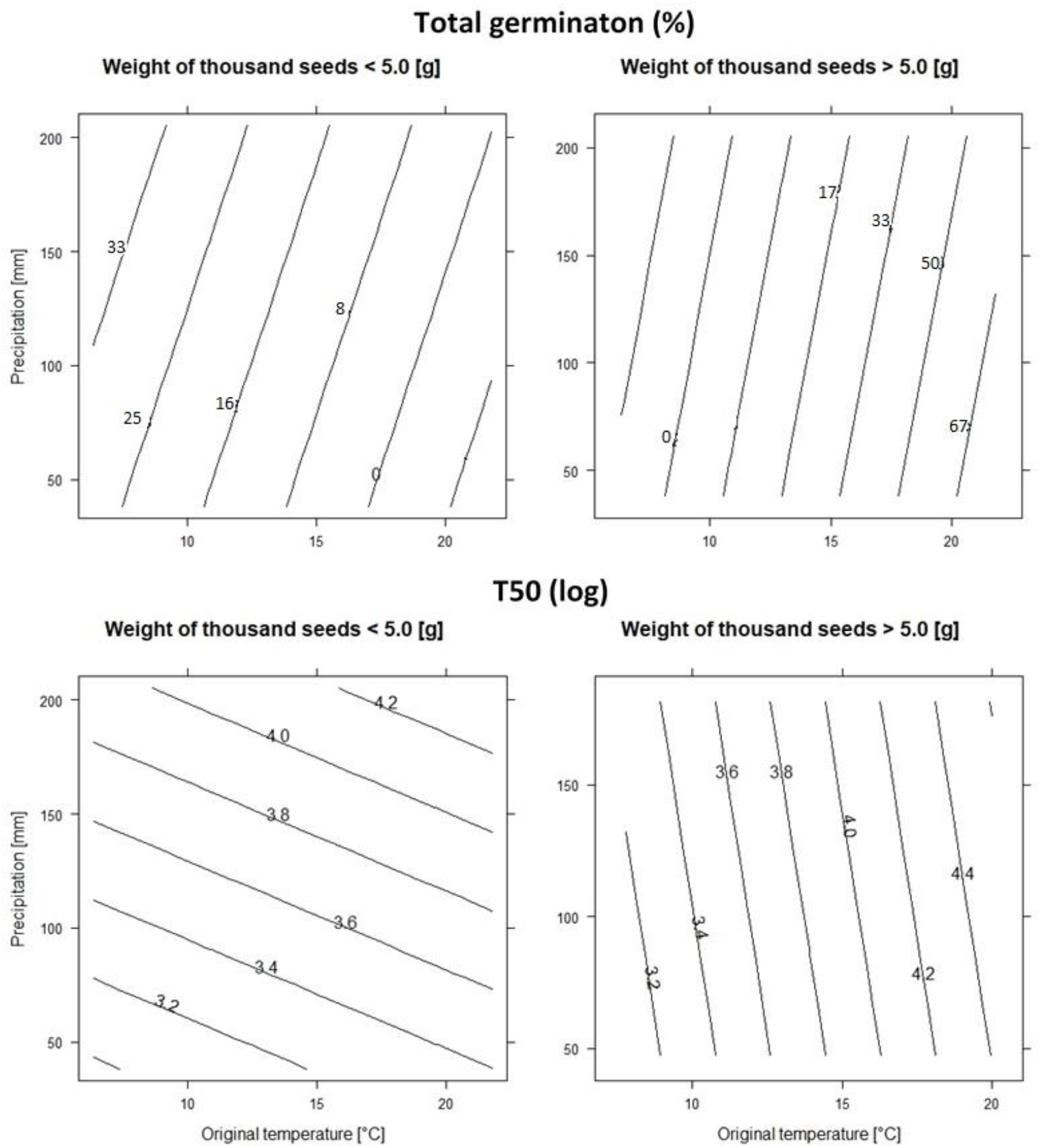
Effect of seed mass, original temperature and original precipitation on total germination (%) and germination speed (T50) (log). For purpose of graphs we divided population according the weight of one thousand seeds to seeds < 5.0 g and ≥ 5.0 g as maximum seed mass was 9.8 g.

**Fig 5.**
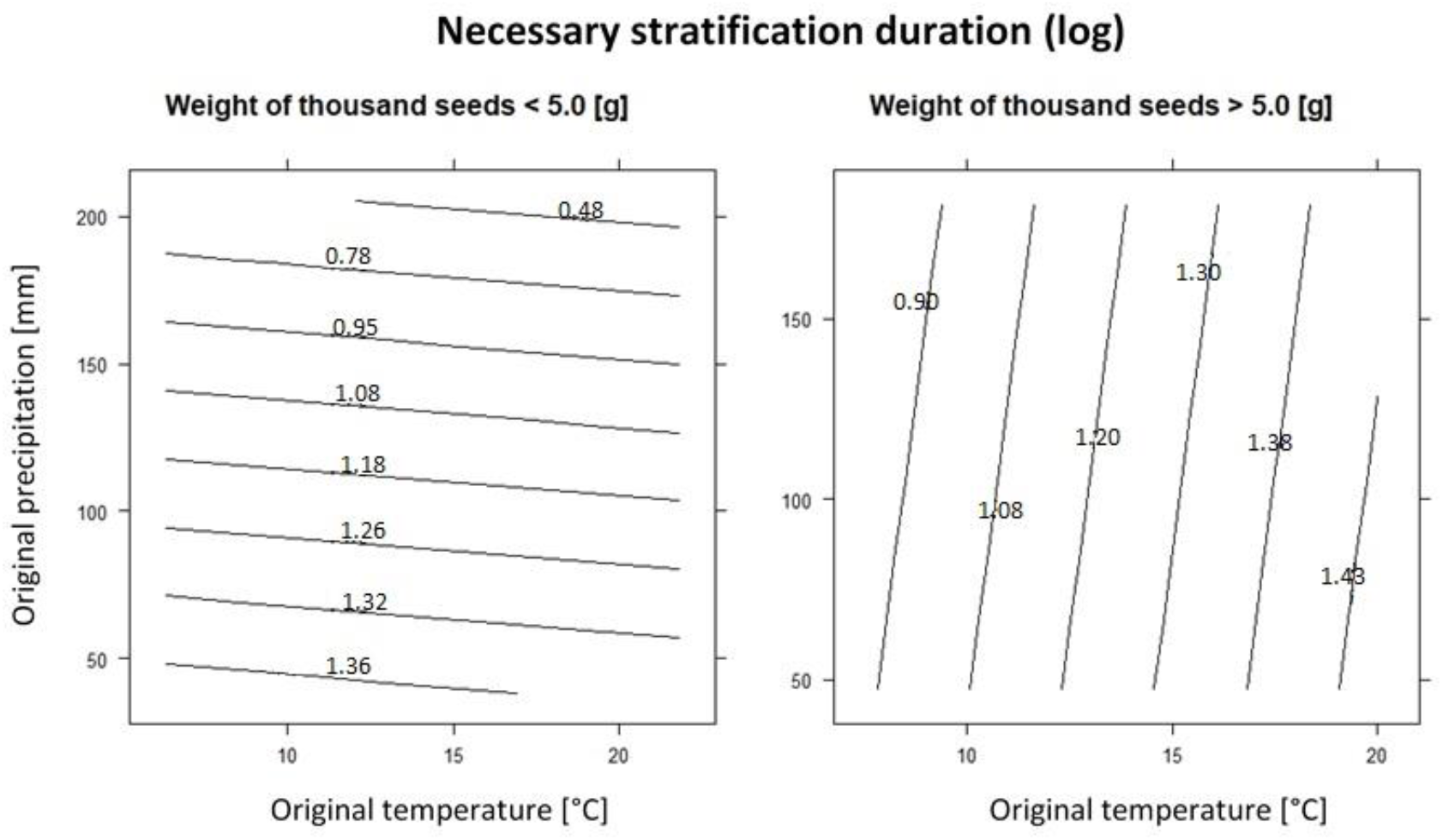
Effect of seeds mass, original temperature and original precipitation on necessary stratification duration in weeks (log). For purpose of the graphs, we divided the populations according to the weight of one thousand seeds between < 5.0 g and > 5.0 g as maximum was 9.8 g.

### Effect of shorter stratification and warm temperature

We found significantly higher total germination (F=3.82, p=0.051) and lower T50 (F=10.06, p=0.003) in seeds kept under stratification until 30% germination and warmer temperature than in seeds exposed to shorter stratification and warmer temperature.

### Effect of phylogeny

Total germination, seed dormancy and necessary stratification duration were not significantly affected by phylogeny (all p-values ≥ 0.091). T50 was significantly affected only by phylogenetic axis 2 (F=6.30, p=0.031) and seed mass was significantly influenced by both phylogenetic axes (F=39.55, p<0.001 and F=16.19, p<0.001, respectively).

All the previously significant predictors of T50 became non-significant after accounting for phylogeny (Table 1). In contrast, the results for seed mass remained unaffected by considering species phylogeny as all the other predictors were not significant already prior to accounting for phylogeny (all p-values ≥ 0.246).

## Discussion

The germination characteristics were affected by seed mass and climate with only limited effects of phylogeny, with the highest variation being attributable to the interaction between original conditions and seed mass. Light seeds were more affected by original precipitation than heavy seeds. Decreasing target temperature increased germination even in seeds from populations in lower altitudes. Shorter stratification duration followed by warmer temperature reduced total germination and increased germination speed (T50). This suggests that climate change could potentially affect both timing of germination and the overall rate of recruitment from seeds. Phylogeny significantly affected only seed mass and germination speed.

### Effect of seed origin, target conditions and seed mass on germination characteristics

Despite a range of previous studies demonstrating that germination depends on original conditions (e.g. (Meyer et al. 1997) (Santo et al. 2015) (Mira, Arnal and Perez-Garcia 2017)), original environment affected only necessary stratification duration in our study. Populations coming from colder localities required longer stratification (similarly in (Navarro and Guitian 2003). In higher altitudes cold period lasts longer than in lowlands and species are adapted to this. However, our results indicate that the necessary stratification duration also depends on precipitation. When precipitation is low, the necessary stratification duration is prolonged as seeds are probably waiting for period with enough moisture. This effect is the strongest in combination of low precipitation and warm temperature as insufficiency of water is more dangerous for seeds and seedling in warm conditions (Baskin and Baskin 2001). The lack of significant main effect of original climate on the other germination characteristics is in line with (Chamorro, Luna and Moreno 2013). In our system, it may be explained by many significant interactions between original and target conditions and original conditions and seed mass as discussed below.

Target temperature influenced all studied germination characteristics as demonstrated previously (e.g.(Grime et al. 1981) (Schütz and Rave 1999) (Gardarin et al. 2011)). However, in contrast to (Gardarin et al. 2011) (Ooi, Auld and Denham 2012) (Walder and Erschbamer 2015), total germination decreased and not increased with increasing target temperature. T50 showed an opposite trend, i.e. seeds germinated faster in warmer conditions, which is in line with (Walder and Erschbamer 2015). However, fast germination does not have to be advantageous in all cases, as strong competition especially for light can occur between seedlings (Baskin, Baskin and Parr 1986). (Perglová et al. 2009) demonstrated that *Impatiens* species are able to form short-term persistent seed bank (sensu Thompson 1993). We found the highest number of dormant seeds in the coldest target temperature. The species probably form short-term persistent seed banks in cold rather than in warm conditions. This result, together with results of germination, indicates that cold conditions are the most suitable for *Impatiens* seedling recruitment.

The effects of original and target conditions interacted in their effect on seed germination. This is in line with our previous results from a different system, Veselá et al. (submitted) and study of (Bauk et al. 2017). Although interaction between target and original conditions can show real species germination response to changing climate (Veselá et al. (submitted), these types of studies are still very rare. Such knowledge is necessary for predicting the effects of climate change on the abundance and distribution of species (Butler, Wheeler and Stabler 2012) (Davila, Tellez and Lira 2013) (Aragon-Gastelum et al. 2017). From our results it seems that *Impatiens* will be probably forced to migrate to higher altitudes. Although the highest total germination was recorded in seeds coming from warm and dry localities almost in all target conditions, total germination under predicted future temperature was the lowest. The highest germination was observed in the coldest target temperature. This result is different from our previous study, Veselá et al. (submitted), suggesting that *Anthoxanthum* species, dominant grass species, will likely profit from the expected climate change when the timing of germination and rain meet.

In line with studies of (Navarro and Guitian 2003), (Wu and Du 2007) and (Münzbergová and Plačková 2010), we found that heavier seeds germinate more than lighter ones. This could be caused by higher content of nutrients (Rees et al. 2001). However, to our best knowledge, our results are the first to show that the effects of seed mass interact with original climatic conditions. Heavy seeds followed the above-mentioned trend that seeds coming from the warm and dry localities have the highest germination and germinate fast. In contrast, light seeds have the highest germination when they come from cold and wet localities. Germination is initiated by sucking of water (Fenner and Thompson 2005) and light seeds probably require higher water content in the soil than heavy seeds. This is obvious from results of germination and necessary stratification duration. Necessary stratification duration is the shortest in seeds coming from the wet localities and stratification duration in light-seeded species is more dependent on precipitation than on temperature. Heavy seeds require only short stratification when coming from wet and cold localities. But temperature is more important determinant of necessary stratification duration compared to precipitation. It seems that future changes mainly in temperature and less in precipitation will be crucial for heavy-seeded species, while changes both in temperature and in precipitation will play a role in recruitment of light-seeded species.

### Effect of shorter stratification and warm temperature

It was previously reported that shortening of stratification can lead to lower germination (Garcia-Fernandez et al. 2015) (Carta et al. 2016b). Simultaneously, it was suggested that warmer climate can bring about higher germination (Gardarin et al. 2011) (Ooi et al. 2012) (Walder and Erschbamer 2015) (Veselá et al. submitted). With ongoing climate change, the plants will probably face combination of shorter stratification and warmer temperatures during vegetation period (IPCC 2014). Our results suggest that such temperature changes will lead to faster but reduced germination. As this is the first study providing such data, more experiments are needed to confirm this. In any case, these results indicate that both changes in temperature and stratification need to be considered to obtain reliable insights into species response to future changing climate.

### Effect of phylogeny

Seed traits, dormancy patterns, and germination responses have ancient origins, and therefore, phylogenetic relationships remain an important part of understanding how they vary (Forbis, Floyd and de Queiroz 2002) (Donohue et al. 2010) (Willis et al. 2014) (Dayrell et al. 2017). In line with previous studies (Zhang et al. 2004) (Moles et al. 2005) (Norden et al. 2009) (Barak et al. 2018), seed mass was strongly affected by phylogenetic relationships. As it was not affected by original environment, seed mass of *Impatiens* seems to be a result of constrains over long-standing evolution of the genus.

Absence of the effect of phylogeny on total germination, seed dormancy and necessary stratification duration indicate that these germination characteristics are driven by environmental conditions rather than by phylogeny. Germination response is probably result of species adaptation to their original conditions. Germination speed was the only germination characteristic related to phylogeny. Due to the existence of phylogenetic niche conservatism (Westoby, Leishman and Lord 1995) (Leishman, Westoby and Jurado 1995), it is not possible to distinguish, if the germination speed of *Impatiens* is a result of environment, or whether phylogenetic constraints are more important in determining these patterns. (Carta et al. 2016a) found phylogenetic signal in germination speed within one genus, however, this subject requires further attention.

### Conclusion

This study demonstrates the importance of including interaction of original and target environment into climate change studies. Simultaneously, it highlights strong dependency of necessary stratification duration on original climate. Predicted climate change represented by shorter stratification period followed by warmer temperature negatively affected species germination. Germination of *Impatiens* species will thus be probably negatively affected by climate change forcing the species to migrate to higher altitudes. Germination response of *Impatiens*, with exception of germination speed (T50), is driven by environmental conditions rather than by phylogeny. The only variable affected by phylogeny was seed mass indicating, that unlike germination behaviour, seed mass is unlikely to change with changing conditions. Heavy seeds germinate the best and the fastest when they come from warm and dry localities, while light seeds germinate the best when they come from cold and wet localities. It seems that future changes both in temperature and precipitation will affect *Impatiens* germination, but the effects will differ between heavy and light-seeded species. As seed mass strongly affected species ability to adapt to their original conditions, considering seed mass is crucial for proper predictions of future germination behaviour of the species. The effects of seed mass on species germination patterns thus need to receive more attention in future studies.

## Acknowledgement

We thank Z. Líblová and M. Lokvencová for help with the germination experiment, Z. Líblová and M. Šurinová for providing their unpublished phylogenetic data and Wojciech Adamowski for help with species identification. The study was supported by project GAČR 17-10280S.

## References

Akiyama S, Ohba H (2016) Studies of Impatiens (Balsaminaceae) of Nepal 3. Impatiens scabrida and Allied Species. Bull Natl Mus Nat Sci Ser B Bot 42:121–130

Ackerly, D. D. & M. J. Donoghue (1995) Phylogeny and ecology reconsidered. Journal of Ecology, 83, 730–733.

Aragon-Gastelum, J. L., E. Badano, L. Yanez-Espinosa, H. M. Ramirez-Tobias, J. P. Rodas-Ortiz, C. Gonzalez-Salvatierra & J. Flores (2017) Seedling survival of three endemic and threatened Mexican cacti under induced climate change. Plant Species Biology, 32, 92–99.

Barak, R. S., T. M. Lichtenberger, A. Wellman-Houde, A. T. Kramer & D. J. Larkin (2018) Cracking the case: Seed traits and phylogeny predict time to germination in prairie restoration species. Ecology and Evolution, 8, 5551–5562.

Baskin, C. C. & J. M. Baskin (1988) Germination ecophysiology of herbaceous plant-species in a temperate region. American Journal of Botany, 75, 286–305.

Baskin, J. M., C. C. Baskin & J. C. Parr (1986) Field emergence on *Lamium amplexicaule* L. and *Lamium purpureum* L. in relation to the annual seed dormancy cycle. Weed Research, 26, 185–190.

Bates, D., M. Machler, B. M. Bolker & S. C. Walker (2015) Fitting Linear Mixed-Effects Models Using lme4. Journal of Statistical Software, 67, 1–48.

Bauk, K., J. Flores, C. Ferrero, R. Perez-Sanchez, M. L. L. Penas & D. E. Gurvich (2017) Germination characteristics of Gymnocalycium monvillei (Cactaceae) along its entire altitudinal range. Botany, 95, 419–428.

Bu, H. Y., G. Z. Du, X. L. Chen, X. L. Xu, K. Liu & S. J. Wen (2008) Community-wide germination strategies in an alpine meadow on the eastern Qinghai-Tibet plateau: phylogenetic and life-history correlates. Plant Ecology, 195, 87–98.

Butler, C. J., E. A. Wheeler & L. B. Stabler (2012) Distribution of the threatened lace hedgehog cactus (Echinocereus reichenbachii) under various climate change scenarios. Journal of the Torrey Botanical Society, 139, 46–55.

Carta, A., S. Hanson & J. V. Muller (2016a) Plant regeneration from seeds responds to phylogenetic relatedness and local adaptation in Mediterranean Romulea (Iridaceae) species. Ecology and Evolution, 6, 4166–4178.

Carta, A., R. Probert, M. Moretti, L. Peruzzi & G. Bedini (2014) Seed dormancy and germination in three Crocus ser. Verni species (Iridaceae): implications for evolution of dormancy within the genus. Plant Biology, 16, 1065–1074.

Carta, A., R. Probert, G. Puglia, L. Peruzzi & G. Bedini (2016b) Local climate explains degree of seed dormancy in *Hypericum elodes* L. (Hypericaceae). Plant Biology, 18, 76–82.

Cavieres, L. A. & M. T. K. Arroyo (2000) Seed germination response to cold stratification period and thermal regime in *Phacelia secunda* (Hydrophyllaceae) - Altitudinal variation in the mediterranean Andes of central Chile. Plant Ecology, 149, 1–8.

Chamorro, D., B. Luna & J. M. Moreno (2013) Germination response to various temperature regimes of four Mediterranean seeder shrubs across a range of altitudes. Plant Ecology, 214, 1431–1441.

Coolbear, P., A. Francis & D. Grierson (1984) The effect of low-tempereture pre-sowing treatment on the germination performance and membrane integrity of artificially aged tomato seeds. Journal of Experimental Botany, 35, 1609–1617.

Cottrell, H. J. (1947) Tetrazolium salt as a seed germination indicator. Nature, 159, 748–748.

Davila, P., O. Tellez & R. Lira (2013) Impact of climate change on the distribution of populations of an endemic Mexican columnar cactus in the Tehuacan-Cuicatlan Valley, Mexico. Plant Biosystems, 147, 376–386.

Dayrell, R. L. C., Q. S. Garcia, D. Negreiros, C. C. Baskin, J. M. Baskin & F. A. O. Silveira (2017) Phylogeny strongly drives seed dormancy and quality in a climatically buffered hotspot for plant endemism. Annals of Botany, 119, 267–277.

Donohue, K., R. R. de Casas, L. Burghardt, K. Kovach & C. G. Willis (2010) Germination, Postgermination Adaptation, and Species Ecological Ranges. Annual Review of Ecology, Evolution, and Systematics, Vol 41, 41, 293–319.

Dreesen, F. E., H. J. De Boeck, I. A. Janssens & I. Nijs (2014) Do successive climate extremes weaken the resistance of plant communities? An experimental study using plant assemblages. Biogeosciences, 11, 109–121.

Esmaeili, M. M., A. Sattarian, A. Bonis & J. B. Bouzille (2009) Ecology of seed dormancy and germination of *Carex divisa* Huds.: Effects of stratification, temperature and salinity. International Journal of Plant Production, 3, 27–40.

Farooq, M., S. M. A. Basra, N. Ahmad & K. Hafeez (2005) Thermal hardening: A new seed vigor enhancement tool in rice. Journal of Integrative Plant Biology, 47, 187–193.

Figueroa, J. A. & J. J. Armesto (2001) Community-wide germination strategies in a temperate rainforest of Southern Chile: ecological and evolutionary correlates. Australian Journal of Botany, 49, 411–425.

Forbis, T. A., S. K. Floyd & A. de Queiroz (2002) The evolution of embryo size in angiosperms and other seed plants: Implications for the evolution of seed dormancy. Evolution, 56, 2112–2125.

Garcia-Fernandez, A., A. Escudero, C. Lara-Romero & J. M. Iriondo (2015) Effects of the duration of cold stratification on early life stages of the Mediterranean alpine plant Silene ciliata. Plant Biology, 17, 344–350.

Gardarin, A., C. Daurr & N. Colbach (2011) Prediction of germination rates of weed species: Relationships between germination speed parameters and species traits. Ecological Modelling, 222, 626–636.

Gimenez-Benavides, L., A. Escudero & F. Perez-Garcia (2005) Seed germination of high mountain Mediterranean species: altitudinal, interpopulation and interannual variability. Ecological Research, 20, 433–444.

Grime, J. P., G. Mason, A. V. Curtis, J. Rodman, S. R. Band, M. A. G. Mowforth, A. M. Neal & S. Shaw (1981) A comparative study of germination characteristics in a local flora. Journal of Ecology, 69, 1017–1059.

Hijmans, R. J., S. E. Cameron, J. L. Parra, P. G. Jones & A. Jarvis (2005) Very high resolution interpolated climate surfaces for global land areas. International Journal of Climatology, 25, 1965–1978.

Hradilová, I., M. Duchoslav, J. Brus, V. Pechanec, M. Hýbl, P. Kopecký, L. Smržová, N. Štefelová, T. Václavek, M. Bariotakis, J. Machalová, K. Hron, S. Pirintsos & P. Smykal (2019) Variation in wild pea (Pisum sativum subsp. elatius) seed dormancy and its relationship to the environment and seed coat traits. Peerj, 7.

Janssens, S. B., E. B. Knox, S. Huysmans, E. F. Smets & V. Merckx (2009) Rapid radiation of Impatiens (Balsaminaceae) during Pliocene and Pleistocene: Result of a global climate change. Molecular Phylogenetics and Evolution, 52, 806–824.

Knapp, A. K., C. Beier, D. D. Briske, A. T. Classen, Y. Luo, M. Reichstein, M. D. Smith, S. D. Smith, J. E. Bell, P. A. Fay, J. L. Heisler, S. W. Leavitt, R. Sherry, B. Smith & E. Weng (2008) Consequences of More Extreme Precipitation Regimes for Terrestrial Ecosystems. Bioscience, 58, 811–821.

Leishman, M. R., M. Westoby & E. Jurado (1995) Correlates of seed size variation - a comparison among 5 temperate floras. Journal of Ecology, 83, 517–529.

Liu, K., J. M. Baskin, C. C. Baskin, H. Y. Bu, G. Z. Du & M. J. Ma (2013) Effect of Diurnal Fluctuating versus Constant Temperatures on Germination of 445 Species from the Eastern Tibet Plateau. Plos One, 8.

Martin, M. C. (1965) An ecological life history of *Geranium maculatum*. American Midland Naturalist, 73, 111-&.

Meyer, S. E. (1992) Habitat correlated variation in Firecracker penstemon (*Penstemon eatonii gray*-Scrophulariaceae) seed germination response. Bulletin of the Torrey Botanical Club, 119, 268–279.

Meyer, S. E., P. S. Allen & J. Beckstead (1997) Seed germination regulation in Bromus tectorum (Poaceae) and its ecological significance. Oikos, 78, 475–485.

Mira, S., A. Arnal & F. Perez-Garcia (2017) Habitat-correlated seed germination and morphology in populations of Phillyrea angustifolia L. (Oleaceae). Seed Science Research, 27, 50–60.

Moles, A. T., D. D. Ackerly, C. O. Webb, J. C. Tweddle, J. B. Dickie & M. Westoby (2005) A brief history of seed size. Science, 307, 576–580.

Münzbergová, Z., V. Hadincová, H. Skálová & V. Vandvik (2017) Genetic differentiation and plasticity interact along temperature and precipitation gradients to determine plant performance under climate change. Journal of Ecology, 105, 1358–1373.

Münzbergová, Z. & I. Plačková (2010) Seed mass and population characteristics interact to determine performance of Scorzonera hispanica under common garden conditions. Flora, 205, 552–559.

Navarro, L. & J. Guitian (2003) Seed germination and seedling survival of two threatened endemic species of the northwest Iberian peninsula. Biological Conservation, 109, 313–320.

Norden, N., M. I. Daws, C. Antoine, M. A. Gonzalez, N. C. Garwood & J. Chave (2009) The relationship between seed mass and mean time to germination for 1037 tree species across five tropical forests. Functional Ecology, 23, 203–210.

Ooi, M. K. J., T. D. Auld & A. J. Denham (2012) Projected soil temperature increase and seed dormancy response along an altitudinal gradient: implications for seed bank persistence under climate change. Plant and Soil, 353, 289–303.

Paulů, A., L. Harčariková & Z. Münzbergová (2017) Are there systematic differences in germination between rare and common species? A case study from central European mountains. Flora, 236–237, 15–24.

Perglová, I., J. Pergl, H. Skalová, L. Moravcová, V. Jarošík & P. Pyšek (2009) Differences in germination and seedling establishment of alien and native *Impatiens* species. Preslia, 81, 357–375.

Press, J.R., K.K. Shrestha & D.A. Sutton (2000) Annotated Checklist of the Flowering Plants of Nepal. London: The Natural History Museum

Qaderi, M. M. & P. B. Cavers (2002) Interpopulation and interyear variation in germination in Scotch thistle, Onopordum acanthium L., grown in a common garden: Genetics vs environment. Plant Ecology, 162, 1–8.

Rees, M., R. Condit, M. Crawley, S. Pacala & D. Tilman (2001) Long-term studies of vegetation dynamics. Science, 293, 650–655.

Santo, A., E. Mattana & G. Baechetta (2015) Inter- and intra-specific variability in seed dormancy loss and germination requirements in the Lavatera triloba aggregate (Malvaceae). Plant Ecology and Evolution, 148, 100–110.

Schütz, W. & P. Milberg (1997) Seed dormancy in *Carex canescens*: Regional differences and ecological consequences. Oikos, 78, 420–428.

Schütz, W. & G. Rave (1999) The effect of cold stratification and light on the seed germination of temperate sedges (Carex) from various habitats and implications for regenerative strategies. Plant Ecology, 144, 215–230.

Seglias, A. E., E. Williams, A. Bilge & A. T. Kramer (2018) Phylogeny and source climate impact seed dormancy and germination of restoration-relevant forb species. Plos One, 13.

Song, Y., Y. M. Yuan & P. Kupfer (2003) Chromosomal evolution in Balsaminaceae, with cytological observations on 45 species from Southeast Asia. Caryologia, 56, 463–481.

Tatebe, H., M. Ishii, T. Mochizuki, Y. Chikamoto, T. T. Sakamoto, Y. Komuro, M. Mori, S. Yasunaka, M. Watanabe, K. Ogochi, T. Suzuki, T. Nishimura & M. Kimoto (2012) The Initialization of the MIROC Climate Models with Hydrographic Data Assimilation for Decadal Prediction. Journal of the Meteorological Society of Japan, 90A, 275–294.

Tingstad, L., S. L. Olsen, K. Klanderud, V. Vandvik & M. Ohlson (2016) Temperature, precipitation and biotic interactions as determinants of tree seedling recruitment across the tree line ecotone (vol 179, pg 599, 2015). Oecologia, 180, 917–918.

Walck, J. L., S. N. Hidayati, K. W. Dixon, K. Thompson & P. Poschlod (2011) Climate change and plant regeneration from seed. Global Change Biology, 17, 2145–2161.

Walder, T. & B. Erschbamer (2015) Temperature and drought drive differences in germination responses between congeneric species along altitudinal gradients. Plant Ecology, 216, 1297–1309.

Wang, J. H., C. C. Baskin, X. L. Cui & G. Z. Du (2009) Effect of phylogeny, life history and habitat correlates on seed germination of 69 arid and semi-arid zone species from northwest China. Evolutionary Ecology, 23, 827–846.

Wang, J. H., W. Chen, C. C. Baskin, J. M. Baskin, X. L. Cui, Y. Zhang, W. Y. Qiang & G. Z. Du (2012) Variation in seed germination of 86 subalpine forest species from the eastern Tibetan Plateau: phylogeny and life-history correlates. Ecological Research, 27, 453–465.

Westoby, M., M. R. Leishman & J. M. Lord (1995) On misinterpreting the phylogenetic correction. Journal of Ecology, 83, 531–534.

Willis, C. G., C. C. Baskin, J. M. Baskin, J. R. Auld, D. L. Venable, J. Cavender-Bares, K. Donohue, R. R. de Casas & N. E. G. W. Grp (2014) The evolution of seed dormancy: environmental cues, evolutionary hubs, and diversification of the seed plants. New Phytologist, 203, 300–309.

Wu, G. L. & G. Z. Du (2007) Germination is related to seed mass in grasses (Poaceae) of the eastern Qinghai-Tibetan Plateau, China. Nordic Journal of Botany, 25, 361–365.

Wu, G. L., W. Li & G. Z. Du (2011) Relationship between germination and seed size in alpine shrubs in Tibetan Plateau. Pakistan Journal of Botany, 43, 2793–2796.

Xu, J., W. L. Li, C. H. Zhang, W. Liu & G. Z. Du (2014) Variation in Seed Germination of 134 Common Species on the Eastern Tibetan Plateau: Phylogenetic, Life History and Environmental Correlates. Plos One, 9.

Yu, S. X., S. B. Janssens, X. Y. Zhu, M. Liden, T. G. Gao & W. Wang (2016) Phylogeny of Impatiens (Balsaminaceae): integrating molecular and morphological evidence into a new classification. Cladistics, 32, 179–197.

Yuan, Y. M., Y. Song, K. Geuten, E. Rahelivololona, S. Wohlhauser, E. Fischer, E. Smets & P. Kupfer (2004) Phylogeny and biogeography of Balsaminaceae inferred from ITS sequences. Taxon, 53, 391–403.

Zhang, S. T., G. Z. Du & J. K. Chen (2004) Seed size in relation to phylogeny, growth form and longevity in a subalpine meadow on the east of the Tibetan Plateau. Folia Geobotanica, 39, 129–142.

