## Supplement material for "Seed mass and plant origin interact to determine species germination patterns"

Supporting information 1. List of species and location of their collection; altitude (m a.s.l.), latitude (N) and longitude (E).

| **Species** | **Altitude** | **N** | **E** |
|---|---|---|---|
| *Impatiens bajurensis* S. Akiyama et H. Ohba | 1 540 | 29.5939 | 80.4573 |
| *Impatiens balsamina* L. | 1 330 | 27.6838 | 85.2836 |
| *Impatiens bicornuta* s.l. Wall. | 2 060 | 27.8141 | 85.3519 |
| *Impatiens bicornuta* s.l. Wall. | 2 158 | 27.8031 | 85.4236 |
| *Impatiens bicornuta* s.l. Wall. | 2 280 | 28.3691 | 83.7311 |
| *Impatiens bicornuta* s.l. Wall. | 2 300 | 27.8120 | 85.3737 |
| *Impatiens bicornuta* s.l. Wall. | 2 075 | 28.3824 | 83.8427 |
| *Impatiens bicornuta* s.l. Wall. | 2 273 | 27.6573 | 85.2191 |
| *Impatiens bicornuta* s.l. Wall. | 2 628 | 28.5292 | 84.3036 |
| *Impatiens bicornuta* s.l. Wall. | 2 480 | 28.5256 | 84.3114 |
| *Impatiens cymbifera* Hook. f. | 2 300 | 27.6315 | 87.2246 |
| *Impatiens cymbifera* Hook. f. | 2 430 | 28.3824 | 83.8427 |
| *Impatiens devendrae* Pusalkar* | 2 728 | 29.8902 | 80.9272 |
| *Impatiens devendrae* Pusalkar* | 2 175 | 29.8627 | 80.9046 |
| *Impatiens discolor* DC. | 2 356 | 27.6586 | 85.2303 |
| *Impatiens insignis* DC. | 1 460 | 27.7550 | 85.2754 |
| *Impatiens insignis* DC. | 1 604 | 27.5896 | 85.3792 |
| *Impatiens puberula* DC. | 2 080 | 27.8141 | 85.3519 |
| *Impatiens racemosa* DC. | 2 440 | 27.6494 | 85.2327 |
| *Impatiens racemosa* DC. | 2 270 | 27.6561 | 85.2290 |
| *Impatiens racemosa* DC. | 2 280 | 27.6560 | 85.2290 |
| *Impatiens racemosa* DC. | 2 525 | 27.6652 | 85.2046 |
| *Impatiens racemosa* DC. | 2 228 | 27.6573 | 85.2191 |

| | | | |
|---|---|---|---|
| *Impatiens racemosa* DC. | 2 100 | 27.8081 | 85.3711 |
| *Impatiens racemosa* DC. | 2 275 | 28.5291 | 84.3189 |
| *Impatiens scabrida* DC.** | 2 180 | 29.2099 | 80.6141 |
| *Impatiens tricornis* Lindl.*** | 3 700 | 28.6305 | 84.4722 |
| *Impatiens tricornis* Lindl.*** | 1 240 | 28.4918 | 83.6508 |
| *Impatiens tricornis* Lindl.*** | 1 160 | 28.3138 | 83.7672 |
| *Impatiens tricornis* Lindl.*** | 2 688 | 28.5523 | 84.2415 |
| *Impatiens scullyi* Hook. f. | 2 880 | 28.5722 | 84.1939 |
| *Impatiens scullyi* Hook. f. | 2 280 | 28.5291 | 84.3189 |
| *Impatiens scullyi* Hook. f. | 2 360 | 28.3978 | 83.7801 |
| *Impatiens scullyi* Hook. f. | 2 688 | 28.5523 | 84.2415 |
| *Impatiens scullyi* Hook. f. | 2 587 | 28.5517 | 84.2723 |
| *Impatiens sulcata* Hook. f. | 2 805 | 28.4022 | 83.7011 |
| *Impatiens sulcata* Hook. f. | 3 580 | 28.6696 | 84.0176 |
| *Impatiens sulcata* Hook. f. | 3 540 | 28.8184 | 83.8490 |
| *Impatiens sulcata* Hook. f. | 3 280 | 28.6230 | 84.1350 |
| *Impatiens falcifer* Hook. f. | 2 499 | 27.5774 | 85.3995 |

* Sensu (Pusalkar and Singh 2010)

** Sensu (Akiyama and Ohba 2016)

*** Sensu (Akiyama and Ohba 2016). Before revision by (Akiyama and Ohba 2016) usually called *I. scabrida.*

Supporting information 2. The courses of the temperatures during the days in the growth chambers. Degrees centigrade indicate minimum and maximum day temperatures.

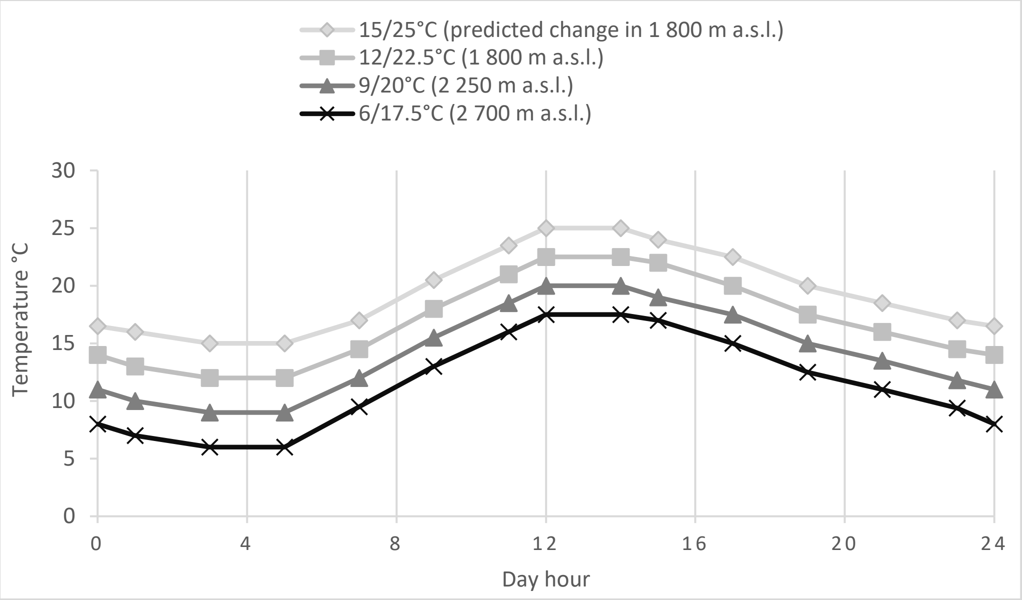

Supporting information 3. Pair correlation matrix showing Pearson correlation coefficients among the dependent variables. Significant values (≤ 0.05) and variables further used for statistical analyses are in bold.

| | **Total germination** | GI | **Seed dormancy** | Seed viability | **T50** | **Necessary stratification duration** |
|---|---|---|---|---|---|---|
| **Total germination** | – | **0.694** | 0.083 | **0.964** | 0.164 | -0.071 |
| GI | | – | -0.288 | **0.577** | **-0.505** | **-0.451** |
| **Seed dormancy** | | | – | **0.345** | **0.626** | **0.466** |
| Seed viability | | | | – | **0.321** | 0.050 |
| **T50** | | | | | – | **0.590** |
| **Necessary strat. duration** | | | | | | – |

Supporting information 4. Effect of original temperature, original precipitation, seed mass and target temperature on total germination, seed dormancy and germination speed (T50) tested in generalized linear mixed effects models with population used as a random factor. Year, longitude, latitude and their interaction were used as covariates in the tests. Results using data including only three target temperature regimes (without 9/20°C) are presented. Significant values (≤ 0.05) are in bold.

| | **Total germination** | | **T50** | | **Seed dormancy** | |
|---|---|---|---|---|---|---|
| | F-value | p-value | F-value | p-value | F-value | p-value |
| Orig.temp | 0.72 | 0.328 | 0.32 | 0.580 | 0.50 | 0.978 |
| Orig.prec | 0.56 | 0.412 | 1.03 | 0.322 | 0.68 | 0.613 |
| Target temp | **5.78** | **0.011** | **3.61** | **0.031** | **5.18** | **0.008** |
| Seed mass | **4.25** | **0.017** | 1.09 | 0.307 | **3.27** | **0.050** |
| Orig.temp:Orig.prec | 0.02 | 0.897 | 0.22 | 0.644 | 0.42 | 0.701 |
| Orig.temp:Target temp | 0.25 | 0.803 | 0.40 | 0.533 | 0.79 | 0.640 |
| Orig.prec:Target temp | **3.91** | **0.044** | **3.53** | **0.033** | 1.07 | 0.154 |
| Orig.temp:Seed mass | **14.89** | **<0.001** | 0.09 | 0.765 | **3.82** | **0.047** |
| Orig.prec:Seed mass | **5.14** | **0.038** | 3.33 | 0.062 | **4.43** | **0.040** |
| Target temp:Seed mass | 0.20 | 0.819 | 0.14 | 0.716 | **4.06** | **0.042** |
| Orig.temp:Orig.prec:Target temp | **12.87** | **<0.001** | 2.17 | 0.154 | 0.21 | 0.591 |
| Orig.temp:Orig.prec:Seed mass | **4.73** | **0.026** | 0.26 | 0.616 | 1.39 | 0.104 |
| Orig.temp:Target temp:Seed mass | 3.11 | 0.075 | 0.11 | 0.747 | 0.30 | 0.744 |
| Orig.prec:Target temp:Seed mass | 0.39 | 0.766 | 2.78 | 0.109 | 2.47 | 0.076 |
| Orig.temp:Orig.prec:Target temp:Seed mass | 0.11 | 0.738 | 1.40 | 0.246 | 2.50 | 0.078 |
